# Self-incompatibility based functional genomics for rapid phenotypic characterization of seed metabolism genes

**DOI:** 10.1101/2024.01.26.577421

**Authors:** Abdul Azeez, Philip D. Bates

## Abstract

Reverse-genetic characterization of plant gene function through technologies such as CRISPR/Cas, RNAi, or gene overexpression requires the ability to efficiently transform the plant species of interest. However, efficient transformation systems are not available for most plant species. *Physaria fendleri* is an oilseed plant valued for its unusual hydroxylated fatty acids (HFA, e.g. lesquerolic acid) that accumulates up to 60% of seed oil and is a non-toxic alternative to castor (*Ricinus communis*) seeds as a source for HFA for the chemical industry. Domestication and improvement of *P. fendleri* seed oil requires characterization of genes involved in developing seed metabolism. Tissue culture-based transformation of *P. fendleri* is laborious, low-efficiency, and time-consuming (T1 ∼18 months). Additionally, *P. fendleri* is self-incompatible requiring laborious hand pollination for propagation and seed collection from transgenic lines. We developed a rapid virus-induced gene silencing (VIGS) method to characterize genes within developing seeds. Identification of the self-incompatibility mechanisms in *P. fendleri* allowed the use of self-compatibility as a novel visual selectable marker by co-targeting the gene of interest (GOI) with the self-incompatibility gene S-locus receptor kinase (SRK). Seeds develop without cross-pollination from silenced SRK and each of those seeds contain the GOI silenced, allowing rapid phenotypic characterization of the seeds in the first generation. Through this methodology we confirmed the in vivo function of two key genes (*FAH12, FAE1*) involved in lesquerolic acid production. Thus, this self-compatibility based functional genomics approach is a rapid methodology for in vivo reverse-genetic gene characterization in self-incompatible plants.

## Introduction

Hydroxylated fatty acids (HFAs) are unusual fatty acids that accumulate in the seed oils of a wide range of plants, with the most abundant accumulation in select species from the *Euphorbiaceae* and *Brassicaceae* plant families (Ohlrogge et al., 2018). Due to the presence of the hydroxyl functional group, HFA can be used in a broader range of chemical applications than the common fatty acids found in food crops, including in the chemical, pharmaceutical, cosmetics, and biofuel industries. The primary industrial source of HFAs is castor (*Ricinus communis*) seed oil, accumulating the HFA ricinoleic acid at ∼80-90% of seed oil. The global castor oil market was USD 1.18 billion in 2018 and is expected to reach USD 1.47 billion by the end of 2025 (Ghosal, 2019). Castor oil, the ricinoleic acid within it, and its chemical derivatives are used in a wide variety of industrial products including: lubricants, soaps, greases, hydraulic fluids, brake fluids, resins, coatings, paints, dyes, inks, biodegradable polyesters, polyurethanes, plastics, nylon-11, waxes, polishes, pharmaceuticals, cosmetics, perfumes, and biofuels (Mutlu and Meier, 2010; Ogunniyi, 2006; Patel et al., 2016). Even though the many uses of HFA from castor oil, castor as a crop has multiple undesirable agronomic and economic features: the seeds contain the highly toxic protein ricin which is one of the most toxic biological agents known (Wilson et al., 1980); castor has limited growth range and adaptability; labor-intensive harvesting; and because castor is mainly grown in semi-arid regions of India without irrigation the seed yield and price is unpredictable due to erratic rainfall patterns (Severino et al., 2012). Therefore, the beneficial and valuable qualities of HFA-containing plant oils, combined with the undesirable features of castor, have led to the desire for alternative sources of HFA that could be grown as a high-value crop.

Multiple species in the *Brassicaceae* family accumulate HFA, are non-toxic, and are adapted to grow in temperate regions of North America (Dierig et al., 1996; Glaser et al., 1992; Ohlrogge et al., 2018). *Physaria fendleri* (previously *Lesquerella fendleri* (Al-Shehbaz and O’Kane, 2002), commonly known as Fendler’s bladderpod) has been considered the prime candidate for domestication, and research toward that end has proceeded over the past several decades such that it is considered an emerging oilseed crop (Brahim et al., 1998; Cruz et al., 2012; Dierig and Ray, 2009; Dierig et al., 1996; Dierig et al., 2011; Harwood et al., 2017; McKeon, 2016; Ploschuk et al., 2005; Qureshi et al., 2018; Salywon et al., 2005; Von Mark and Dierig, 2015; Von Mark et al., 2013). *P. fendleri* accumulates the HFA lesquerolic acid at ∼60% of the seed oil (Brahim et al., 1996; Chen et al., 2009; Cocuron et al., 2014; Lin and Chen, 2013; Lin and Chen, 2014; Lin et al., 2015; Smith Jr et al., 1961). Due to the similar chemical properties of ricinoleic acid and lesquerolic acid, the *P. fendleri* seed oil can be used for many of the same applications as castor oil (Glaser et al., 1992). There is no full genome sequence for *P. fendleri*, but the recent development of molecular markers for breeding (Von Mark et al., 2013), developing seed transcriptomic resources (Horn et al., 2016; Kim and Chen, 2015), and the ability to genetically transform *P. fendleri* (Azeez et al., 2022; Chen, 2011; Chen et al., 2021; Chen et al., 2011; Chen et al., 2016) are aiding crop development. Recently, multiple isotopic tracing studies have indicated that *P. fendleri* developing seeds utilize both nonconventional and novel pathways to partition carbon into fatty acid biosynthesis and assemble oil containing HFA (Bhandari and Bates, 2021; Cocuron and Alonso, 2023). To further understand these novel metabolic pathways and improve the *P. fendleri* seed yield, oil yield and HFA content, we need to understand the *in-planta* function of genes involved in these processes. Despite a protocol for *P. fendleri* tissue culture-based transformation to generate stable transgenic plants, this method suffers from challenges with chimeric tissues and self-incompatibility (endogenous to *P. fendleri*) making production of homozygous knockdown lines laborious, low-efficiency, and time-consuming with ∼18 months as the average time to produce a homozygous transgenic *P. fendleri* line.

Self-incompatibility is a major limitation to improving *P. fendleri* as a high-yielding source of industrially valuable HFA because pollination depends on insect pollinators which can limit total possible seed production. For example, the combination of self-compatible flowers and the presence of pollinators increases crop yield in the related Brassicaceae crop canola (Adamidis et al., 2019). Self-incompatibility is also a hurdle for research progress by making it laborious to produce homozygous transgenic/mutant lines (which requires hand pollination at immature flower stages). It has been reported in several plants that two S-locus genes are responsible for self-incompatibility, the S-locus receptor kinase (SRK) in the stigma, and the S-locus cysteine rich protein (SCR) in the pollen (Kakeda et al., 2008; Nasrallah et al., 2002; Nasrallah et al., 2004; Rahman et al., 2007). To understand and control the undesirable self-incompatibility in *P. fendleri* we adapted virus-induced gene silencing (VIGS) knockdown methods to *P. fendleri* to evaluate self-incompatibility genes. VIGS is a simple, fast, and powerful reverse genetic tool widely used to study plant gene functions (Burch-Smith et al., 2004; Robertson, 2004; Zhirnov et al., 2015). We demonstrate that knockdown of SRK alone is sufficient to induce self-compatibility and produce seed pods without cross pollination. Additionally, we demonstrate at SRK silencing can be utilized as a reporter gene for the co-silencing of developing seed metabolism genes.

VIGS can travel throughout a plant allowing silencing of genes in different tissues (e.g. for evaluation of seed lipid metabolism). However, the transfer of VIGS from an infected plant into developing seed tissue is unpredictable, requiring extensive screening of many seeds to select the seeds containing a silenced gene of interest (GOI). Additionally, due to the self-incompatibility of *P. fendleri*, the production of seed pods of VIGS infected plants first requires laborious forced selfing of immature flowers. The identification of self-incompatibility genes and their efficient silencing by VIGS *fendleri* allowed us to use co-silencing of self-incompatibility as a novel selectable marker for selection of developing seeds harbouring a VIGS construct silencing SRK and a co-targeted seed metabolism GOI. The SRK-GOI silenced seeds can be rapidly identified by visual selection of the developing self-compatible seed pods for further phenotypic characterization of the silenced seed metabolism GOI. Through this methodology we report the *in vivo* confirmation of two key genes (*fatty acid hydroxylase 12, FAH12*, and *fatty acid elongase 1, FAE1*) involved in synthesis of lesquerolic acid in developing *P. fendleri* seeds. Thus, we developed a rapid functional genomics methodology not limited to *P. fendleri* but suitable for a wide range of self-incompatible commercially important crops.

## Results

### Optimization of VIGS infection procedure for *P. fendleri*

The tobacco rattle virus (TRV) spreads quickly throughout the plant with minor symptoms compared to other plant viruses, thus the TRV-mediated VIGS system is most widely used to study plant gene functions (Burch-Smith et al., 2004). In initial testing of *P. fendleri* at the four leaf stage we found low silencing efficiency (mild photobleaching) while targeting *P. fendleri* phytoene desaturase (*PDS*) as a marker by the standard leaf-agroinfiltration method (Burch-Smith et al., 2004). These results suggested that either the infiltration method or transport of the TRV throughout the plant may not work well with mature *P. fendleri* leaf tissues (Figure S1). To improve the silencing efficiency at whole plant level, we utilized vacuum-agroinfiltration of germinated seeds (Figure 1), a method successfully utilized for whole plant level gene silencing in maize and wheat (Zhang et al., 2017). Through parameter optimization (inoculum density, incubation time, etc.) we found 96% silencing efficiency at an agrobacterium OD of 0.4 and 5 min of vacuum-agroinfiltration time with *P. fendleri* germinated seeds (Figure 2a). Among the 96% of plants with photobleaching phenotypes, 20% were completely photobleached (whole plant silencing), whereas 80% had partial photobleaching phenotypes (Figure 2b). Further, we checked the heritability of silencing phenotype, and found around 10% of the seeds showed a photobleached phenotype in the next generation (Figure S2). This result indicates that VIGS-mediated silencing using germinated seeds is not only heritable but also has a silencing effect within seeds, which is our major tissue of interest for evaluation of seed lipid metabolism-related target genes.

**Figure 1:**
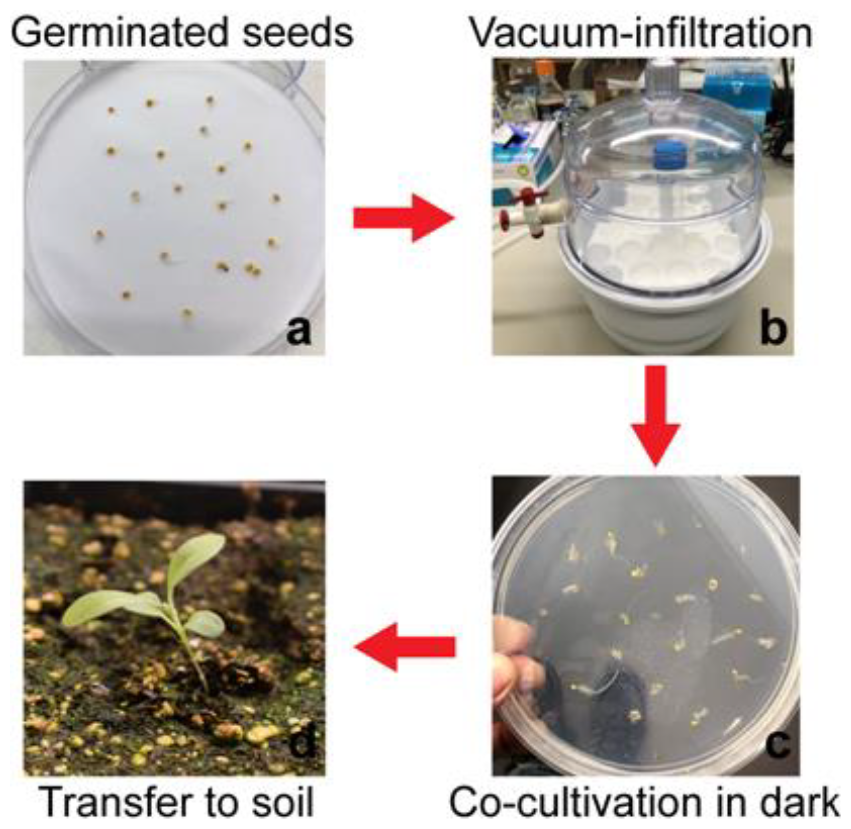
Agrobacterium-mediated vacuum-infiltration of *P. fendleri* (germinated) seeds to Virus Induced Gene Silencing of target genes. (a) 2mM GA3 treated *P. fendleri* seeds to initiate uniform germination. (b) Vacuum-infiltration of germinated seeds with *A. tumefaciens* cells contain VIGS construct of gene of interest. (c) Dry the access agro-solution on filter paper and co-cultivate in dark for one day. (d) Transfer seeds to the soil pots and grow then in growth room.

**Figure 2.**
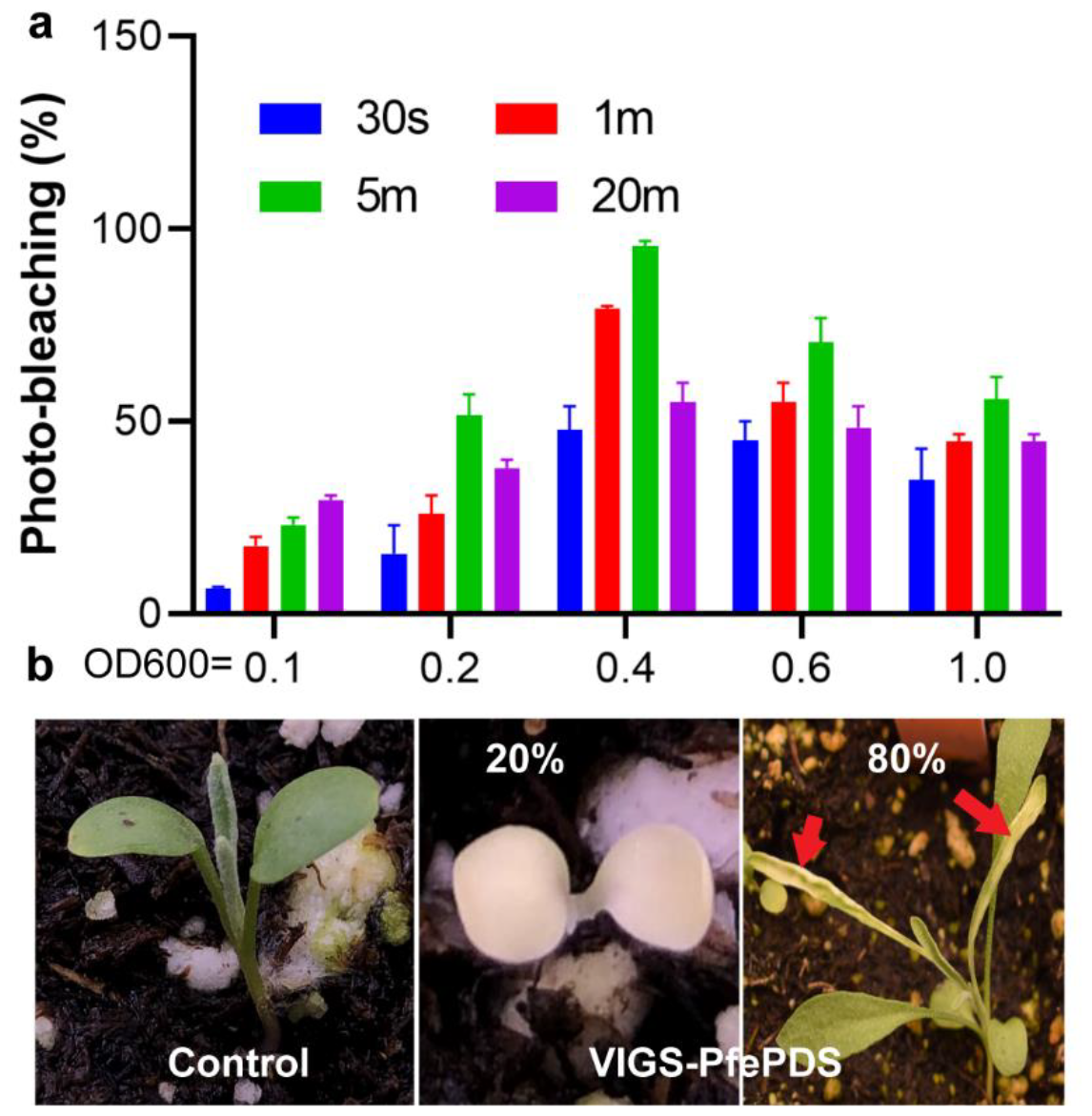
Optimization of silencing efficiency by changing *Agrobacterium* densities and vacuum-infiltration time. (a) 96% silencing efficiency at 0.4 O.D. for 5 min vacuum-infiltration. (b) 20% complete Photobleaching phenotype and 80% with partial Photobleaching phenotype of pTRV2-PfePDS plants.

### VIGS silencing of seed lipid metabolic network genes

To test the VIGS silencing efficiency in on developing *P. fendleri* seed lipid metabolism genes, we selected two genes involved in HFA biosynthesis: the fatty acid hydroxylase 12 (*FAH12* (van de Loo et al., 1995)) and fatty acid elongase 1 (*FAE1* (Kunst et al., 1992; Rossak et al., 2001)). FAH12 hydroxylates oleic acid (18:1) on membrane lipid phosphatidylcholine (PC) producing ricinoleic acid (18:1-OH). After removal of ricinoleic acid from PC and esterification to Coenzyme A, FAE1 is involved in the elongation of ricinoleic acid by two carbons producing lesquerolic acid (20:1-OH) (Chen et al., 2021) esterified to CoA that is then utilized for oil biosynthesis. The knockdown of the FAH12 should limit the conversion of 18:1 to 18:1-OH, and thus 20:1-OH accumulation. The *FAH12_VIGS* inoculation on germinating *P. fendleri* seeds which were then grown to maturity produced seeds with increased 18:1 and reduced 20:1-OH as expected with reduced FAH12 activity (Figure 3a). *FAE1* silencing should limit the elongation of 18:1-OH to produce 20:1-OH. The *FAE1_VIGS* line produced seeds with higher 18:1-OH and less accumulation of the 20:1-OH as expected (Figure 3b). Thus, we have demonstrated that a VIGS system employing germinated seed transformation can be utilized for developing seed gene functional evaluation. Additionally, it is a rapid (only one plant generation, e.g. 4 months as opposed to 18 months of tissue culture-based RNAi transformation) and a less laborious tool than published tissue culture-based methods (Azeez et al., 2022; Chen, 2011; Chen et al., 2021). Although, two drawbacks remained: (1) due to the nature of *P. fendleri* self-incompatibility, laborious hand pollination at immature flower stages is needed for the VIGS inoculated plants to produce seeds; (2) due to the variability of TRV transport throughout the plant and into germline tissue, many individual seeds needed to be screened for fatty acid content to evaluate the metabolic effect of the gene silencing. For example, identification of the top lines in Fig. 3 was found after lipid screening seeds from 40-60 pollinated flowers per line. Thus, self-incompatibility and lack of a marker for which seeds contain the VIGS construct are time and labor consuming limitations to producing a high-throughput VIGS system for the analysis of developing *P. fendleri* seed metabolism. To further develop the VIGS system we investigated the mechanisms of self-incompatibility in *P. fendleri*.

**Figure 3.**
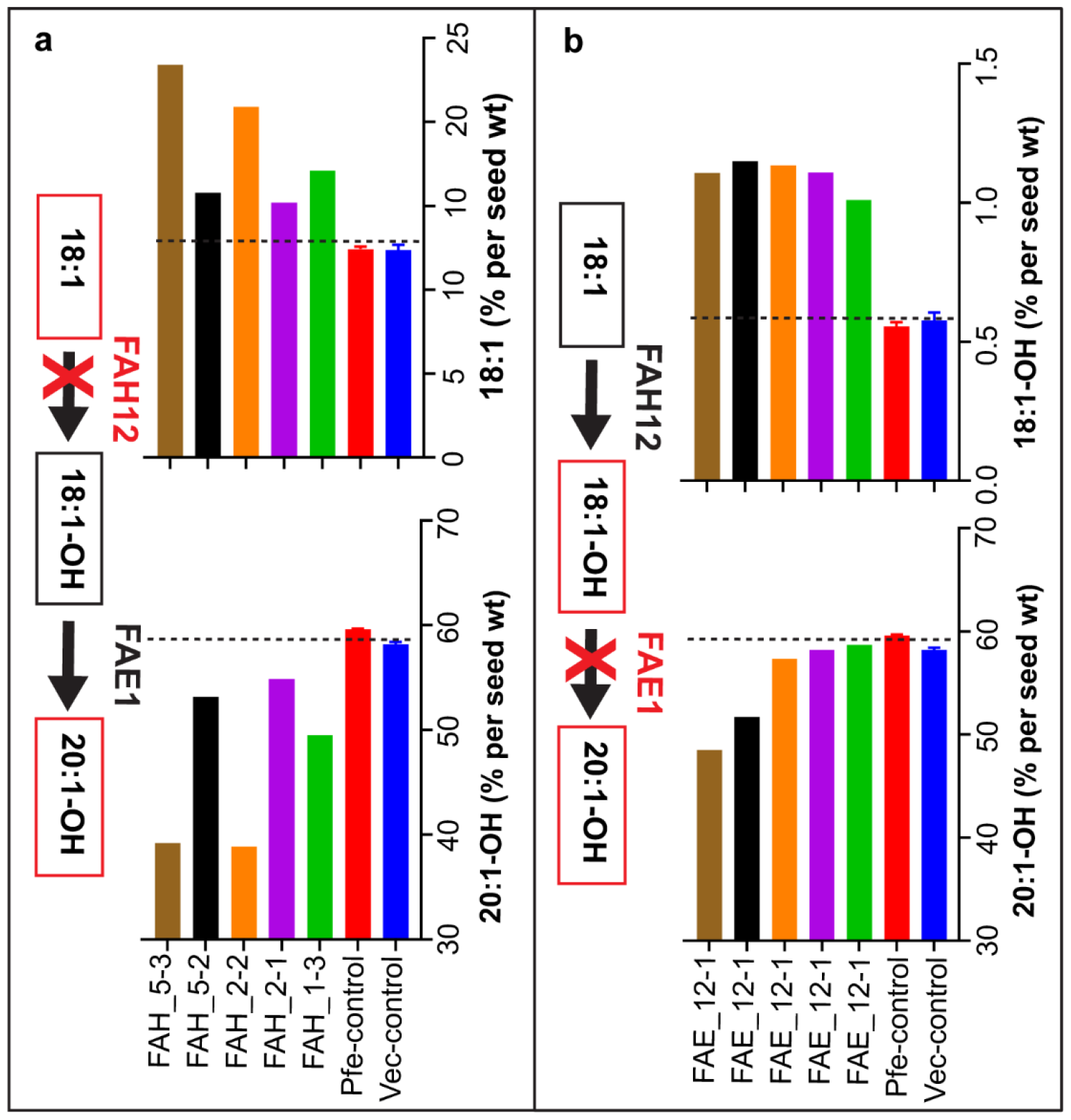
Fatty acid composition in *P. fendleri* FAH12 and FAE1 VIGS seeds. (a) More substrate (18:1) accumulation and decrease in downstream product (20:1-OH) in FAH12-VIGS lines. (b) More substrate 18:1-OH and decrease in product (20:1-OH) in FAE1-VIGS corresponding lines. Amount of FAs calculated based on weight percent of total fatty acids. Two controls used for analysis, empty vector (Vec-control) agro-infiltrated seeds and wild-type *P. fendleri* seeds (Pfe-control), data represents mean ± SEM (*n*= 3), whereas seeds from individual pods were used for VIGS seeds analysis.

### *P. fendleri* S-locus receptor kinase (SRK) is responsible for self-incompatibility

In various Brassicaceae members the S-locus receptor kinase (SRK) and S-locus cysteine-rich (SCR) proteins are the major players inducing self-incompatibility (Nasrallah et al., 2002; Nasrallah et al., 2004). The female determinant SRK is expressed in the stigma and acts as a receptor for pollen-expressed SCR ligand protein. When self-pollen land on the stigma, the SCR ligand binds to the SRK receptor and activate downstream signaling; resulting in rejection of self-pollen (Nasrallah et al., 2002; Nasrallah et al., 2004). *P. fendleri* is a closely related Brassicaceae species to Arabidopsis, therefore we evaluated the role of these two crucial genes in *P. fendleri* self-incompatibility. Similar to other Brassicaceae *P. fendleri* female determinant SRK was expressed primarily in the stigma and male determinant SCR in pollen (Figure 4a). Interestingly, *P. fendleri* is self-compatible at an early stage (before the flower opens) (Chen et al., 2011). The *SRK* expression was low at very early stage 1 and then highly upregulated at stage 2 consistent with our ability to produce homozygous plants through hand pollination only with immature stage 1 flowers (Figure 4b,c). Later, in stages 3 and 4, the *SRK* expression level is reduced (Figure 4c), but stigmas are less receptive to self-pollen than in earlier stages likely due to protein produced during stage 2 that remains in the stigma. To functionally characterize the role of these genes in *P. fendleri* self-incompatibility, we prepared VIGS constructs to knockdown SRK and SCR genes *in planta*. The VIGS-SCR lines produced no seed pods after self-pollination, similar to control plants (Figure 5a, Figure S3). However, ∼30% of the VIGS-SRK lines produced healthy seed pods after self-pollination (Figure 5b). These results indicate that female determinant SRK is the key player in controlling the self-incompatibility of *P. fendleri*.

**Figure 4.**
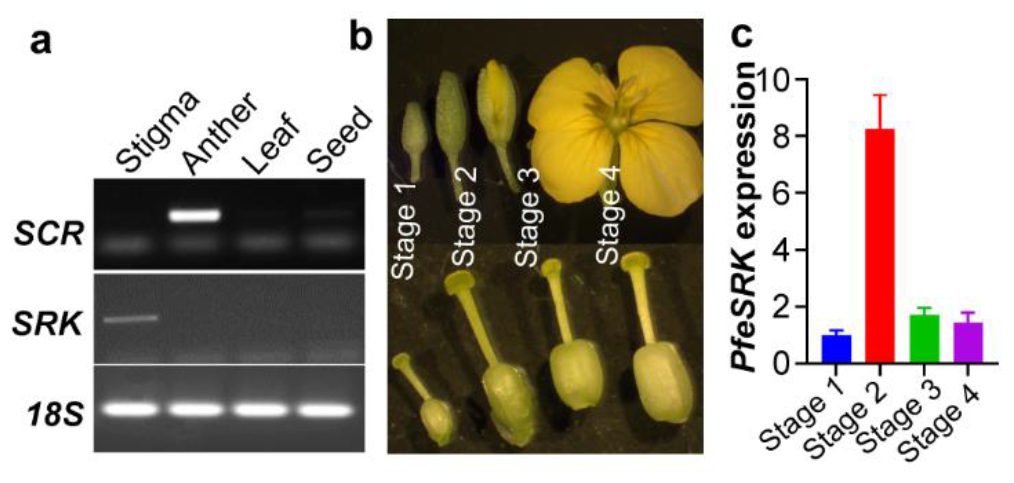
Spatial and temporal expression of *P. fendleri* self-incompatibility genes. (a) *P. fendleri* self-incompatibility (SI) gene S-locus cysteine-rich (*PfeSCR)* specifically expressed in anther whereas S-locus receptor kinase (*PfeSRK)* gene expressed in stigma, semi-quantitative PCR, 18S used as internal control. (b) Developmental stages of *P. fendleri* flower and respective carpel (c) Relative expression of the SRK gene in different development stages of stigma. Reference gene 18S expression used to normalize the target gene expression and expression values are the average of three biological replicates ± SE, and each stage represent pool of 10 stigmas. List of the primers used for PCR is presented in Supplementary Table S1.

**Figure 5.**
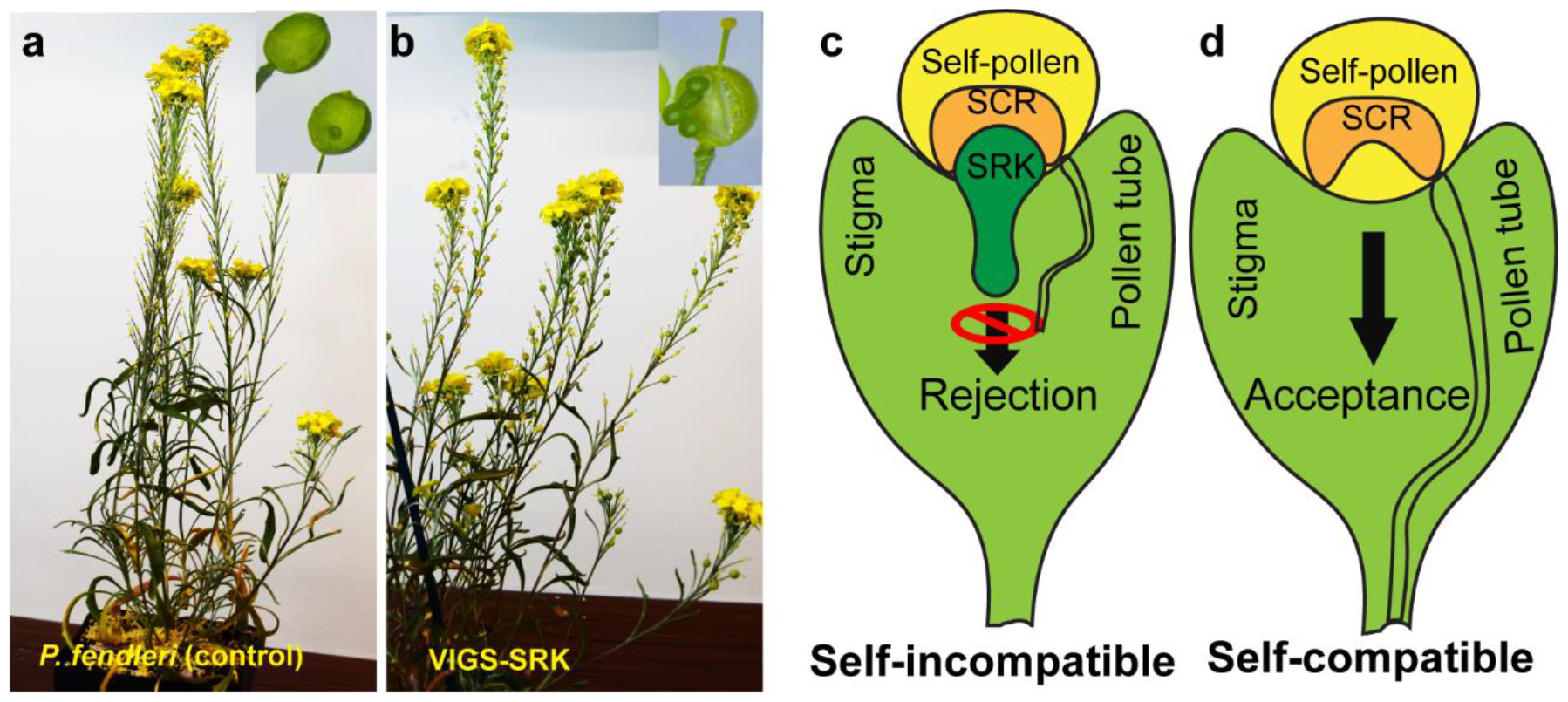
Hypothetical model to generate self-compatible *P. fendleri*, and result of VIGS of self-incompatibility genes. (a) Self-incompatible *P. fendleri* produced no seed after self-pollination. (b) 30% of *P. fendleri* plants produced seed after Virus-induced gene silencing of SRK. (c, d) Self-incompatibility model, self-pollen containing functional SCR protein bind to active SRK expressed in stigma to activate downstream signaling to reject self-pollen. SRK knock-out in *P. fendleri*, in the absence of functional SRK, the stigma will be able to accept self-pollen and grow the pollen tube and eventually plant could become self-compatible.

### Rapid and efficient functional genomics tool for characterization of developing seed metabolic genes

The identification of the genetic factors controlling self-incompatibility in *P. fendleri* and its successful silencing by VIGS (Figure 5) allowed us to propose utilizing silencing of self-incompatibility to both reduce the labor involved in *P. fendleri* VIGS approaches and utilize it as a novel selectable marker. By co-targeting of SRK and our gene of interest (GOI) in a single VIGS construct we can eliminate the need for hand pollination and select just the seed pods containing the silenced genes for further analysis by visual selection of developing pods (Figure 6a, b). In this way we can overcome both of the time-consuming and labor-intensive issues that remained in our prior VIGS methodology.

**Figure 6.**
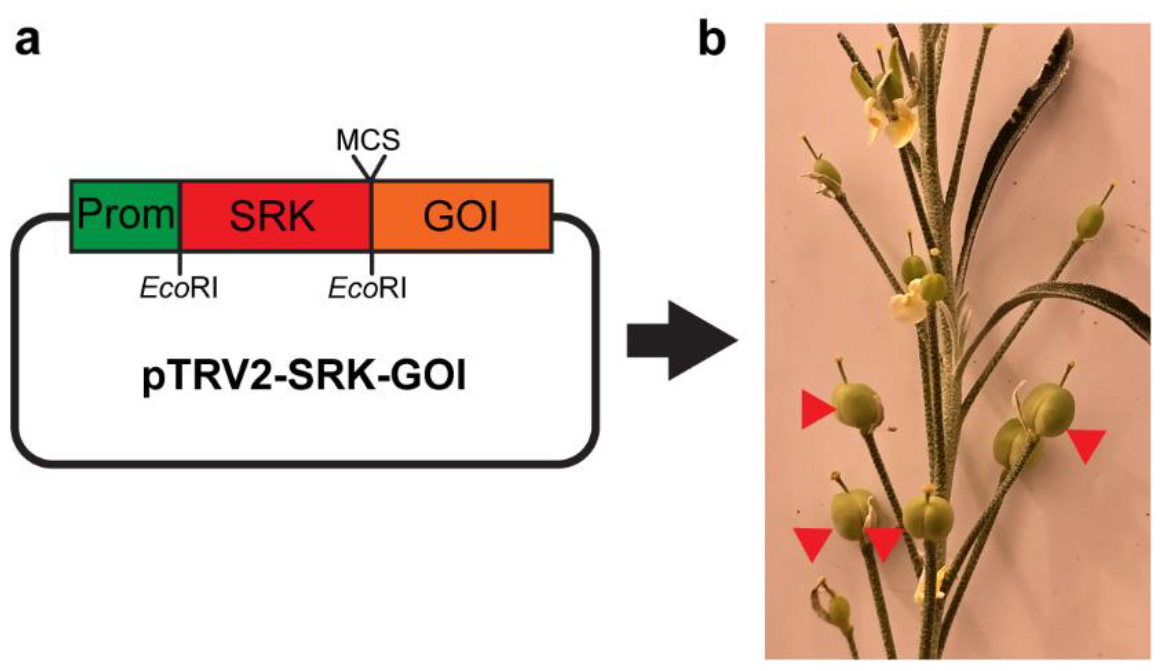
Self-complimentary based a rapid functional genomics tool with visual selection. (a) Cloning strategy of gene of interest (GOI) co-targeted with self-incompatibility gene SRK. (b) Visual selection of the seed pods by co-targeting SRK with GOI.

To test our self-incompatibility based functional genomics tool, we silence FAE1 by co-targeting SRK. We generated several SRK-FAE1 VIGS lines using our protocol, and around 20% of SRK-FAE1 VIGS lines successfully developed seed pods without outcrossing. Further, we confirmed the SRK suppression using qRT-PCR, and found that all the developing seeds produced by the VIGS-SRK-FAE1 and VIGS-SRK control lines had decreased SRK expression as compared to the *P. fendleri* control plants (Figure 7a). The VIGS-SRK-FAE1 mature seeds accumulated more 18:1-OH and less 20:1-OH similar to the VIGS-FAE1 alone (Fig. 3b), confirming suppression of FAE1 in the seeds that developed in without cross pollination (Figure 7b, c).

**Figure 7.**
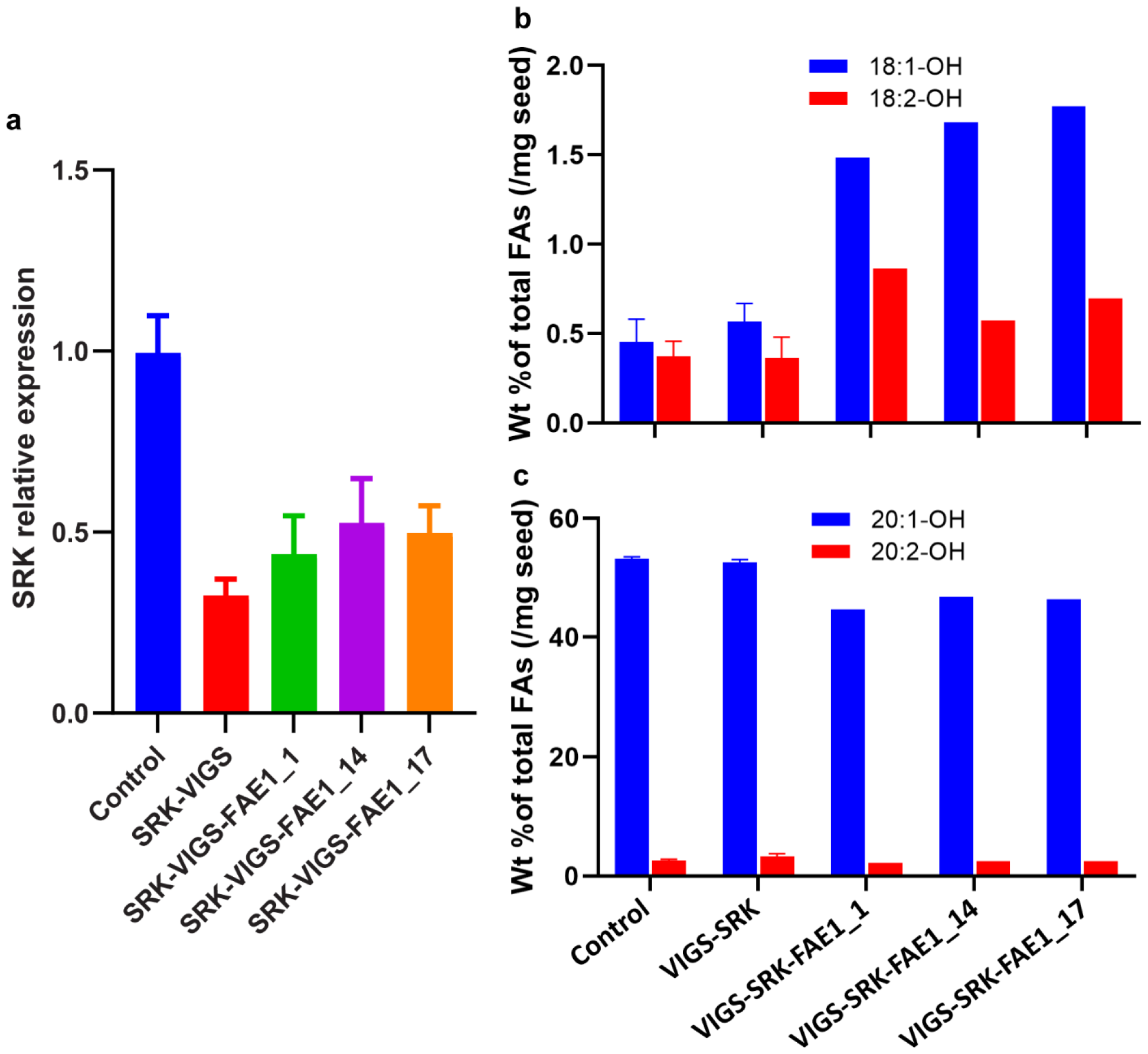
Fatty acid composition in *P. fendleri* VIGS-SRK-FAE1 seeds. (a) Relative expression of the *PfeSRK* gene in flower stigmas of VIGS-SRK-FAE1 lines, expression quantified by qRT-PCR and normalized to the reference *18S* gene. Expression values are the average of three technical replicates ± SE, and each line represent pool of 15 stigmas. (b) More substrate (18:1-OH, 18:2-OH) accumulation and (c) decreased product (20:1-OH, 20:2-OH) in VIGS-SRK-FAE1 seeds. Amount of FAs calculated based on weight percent of total fatty acids. Two controls used for analysis, normal *P. fendleri* seeds (control), and VIGS-SRK seeds and data represents mean ± SEM (*n*= 3), whereas 5 seeds were used for VIGS-SRK-FAE1 for analysis.

Therefore, by combining VIGS of SRK with a GOI in a single construct we developed a novel tool for visual selection of GOI silenced seed pods (those that grow without cross-pollination), thus reducing both the pollination and seed phenotyping labor required to screen various genes for their role in *P. fendleri* seed development and seed metabolic networks such as lipid metabolism. This technology can be replicated in other commercially important self-incompatible crops by co-targeting the GOI with the endogenous SRK, or other known self-incompatibility factors within that species (Fujii et al., 2016).

## Discussion

### VIGS system for developing seed gene functional evaluation

Although *P. fendleri* was discovered in the 1960s for its unique unusual HFA fatty acids, lack of knowledge on how *P. fendleri* accumulate the valuable HFA and its endogenous genetic self-incompatibility have limited the ability to improve *P. fendleri* as a crop through bioengineering. We optimized a rapid VIGS system for *P. fendleri* using vacuum-agroinfiltration of germinated seeds by means of a well-studied marker gene phytoene desaturase (*PfePDS*) and confirmed its function in *P. fendleri* (Figure 1, 2). Using our optimized VIGS system, we characterized *P. fendleri* self-incompatibility genes and demonstrated that the SRK is the key player in controlling self-incompatibility within *P. fendleri* (Figure 4, 5). The application of our VIGS procedure to evaluate developing seed lipid metabolism confirmed *in planta* within *P. fendleri* the roles of FAH12 and FAE1 within lesquerolic acid production (Figure 3, 7). The FAE1 results are consistent with prior results utilizing more labor-intensive stabile transformation of *FAE1_RNAi* in *P. fendleri* (Chen et al., 2021). Additionally, the *P. fendleri* FAH12 activity was previously characterized through heterologous expression of in Arabidopsis (Broun et al., 1998), here we provide the first in vivo results within *P. fendleri* confirming the role of *PfeFAH12* in HFA production. By targeting *P. fendleri* FAH12 and FAE1 to the best of our knowledge this is the first time that the VIGS system has been utilized for gene functional evaluation of developing seed lipid metabolism.

### Rapid functional genomics tool

Self-incompatible plants like *P. fendleri* do not produce seed unless cross pollinated from a different individual of the same species. However, the ability to induce self-compatibility by VIGS allows us to use self-incompatibility as selection tool. We developed a rapid and efficient self-incompatibility-based functional genomics technology to characterize genes involved in seed development that solves two of the major labor-intensive aspects of seed gene characterization through VIGS: (1) the required hand pollination with immature flowers; and (2) the need to screen many seed pods for phenotypes due to the variability of VIGS transfer from the mother plant. In our method by co-targeting the silencing of a seed metabolism GOI with the self-incompatibility gene SRK, hand cross pollination with immature flowers is not required and it produces a rapid visual selection for identifying the seed pods that contain the silenced genes.

Self-incompatibility is a mechanism to improve genetic diversity and is widespread among plants. More than 100 plant families are self-incompatible, which is approximately 40% of the flowering plant species (Igic et al., 2008). Molecular regulation of self-incompatibility systems of three families Brassicaceae, Papaveraceae and Solanaceae are well characterized (Fujii et al., 2016; Kitashiba and Nasrallah, 2014). The Brassicaceae and Papaveraceae self-incompatibility systems are based on self-specific interaction between the male and female S-determinates to reject self-pollen, where as in Solanaceae self-pollen are rejected in the absence of self-recognition of the pollen on the stigma (Fujii et al., 2016). S-RNase ribonuclease activity is also critical for self-pollen rejection in *Petunia inflata* (Huang et al., 1994) and citrus (Hu et al., 2023; Liang et al., 2020). Therefore, our self-incapability based functional genomics technology can be replicated in other commercially important self-incompatible crops containing systems of self-specific molecular interactions by co-targeting GOI with the self-incompatibility factors in those species.

### New tools indicate understanding of unconventional and novel metabolism in *P. fendleri* is on the horizon that can lead to enhancement of *P. fendleri* as a crop

Recently, a novel pathway of triacylglycerol remodeling was proposed for assembly of lesquerolic acid containing oils in developing *P. fendleri* seeds based on in vivo isotopic tracing (Bhandari and Bates, 2021), yet the genes involved have yet to be identified. Likewise, metabolic flux analysis identified *P. fendleri* embryos to be less efficient at metabolizing carbon into biomass than other oilseeds, and that *P. fendleri* utilizes non-conventional pathways (mainly the Rubisco shunt) to channel carbon into oil biosynthesis (Cocuron and Alonso, 2023). The in vivo confirmation of the genes involved in these pathways will provide targets for crop improvement through gene editing and/or biotechnology, the methodology presented here can provide a rapid means to gain further incite on the control of *P. fendleri* metabolism.

The in vivo gene confirmation results presented here have already provided potential targets to enhance *P. fendleri* as a valuable replacement to castor as a source of HFA. For example, confirmation of the role of FAE1 in elongation of ricinoleic acid to lesquerolic acid may indicate that gene editing of FAE1 could be used to produce ricinoleic acid in *P. fendleri* which would be equivalent to castor oil. Additionally, by identifying the controlling factor of self-incompatibility to be SRK, gene editing of SRK has the possibility to produce fully self-compatible *P. fendleri* which may enhanced seed yield when combined with pollinators as in Brassicaceae crop canola (Adamidis et al., 2019). Together, this rapid self-incompatibility based functional genomics approach will be a valuable tool for understanding *P. fendleri* metabolism and seed development for crop improvement, as well as in other self-incompatible plants.

### Experimental procedures

#### Constructs preparation

All primers used were designed based on genomic DNA sequences from the NCBI database (https://blast.ncbi.nlm.nih.gov/Blast.cgi?PAGE_TYPE=BlastSearch&PROG_DEF=blastn&BLAST_SPEC=Assembly&UID=99421664) and purchased from Sigma. Supplementary Table S1 contains primer sequences. For the construction of pTRV2-PfePDS, a fragment of 872 bp of *P. fendleri PDS* cDNA was amplified using PfePDS_F and PfePDS_R primers and cloned into *Eco*RI-*Xba*I sites of pTRV2 vector. To prepare pTRV2-PfeFAE1 and pTRV2-PfeFAH12 constructs a fragments of 591 bp and 423 bp were amplified by respective primers using seed cDNA as templated and cloned at *Kpn*I-*Sac*I and EcoRI-BamHI sites of pTRV2 respectively. For the construction of pTRV2-PfeSRK and pTRV2-PfeSCR vectors, fragments of 543 bp and 251 bp were amplified using *P. fendleri* stigma and anthers cDNA as templates, respectively. PCR products were purified and cloned in pTRV2 vector at *Eco*RI-*Bam*HI and *Bam*HI-*Kpn*I sites, respectively. To prepare the dual target construct of pTRV2-SRK-FAE1, *FAE1* fragment was cloned at *Bam*HI-*Sac*I site of pTRV2-SRK vector.

### Expressional analysis of *P. fendleri* self-incompatibility genes

*P. fendleri* flower stigmas, anthers, leaf, and seed were harvested and immediately frozen in liquid nitrogen and stored at -80°C till used for RNA extraction. Total RNA was extracted using a Spectrum plant total RNA kit (Sigma). Total RNA (10 μg) was treated with RNase-Free DNase (Qiagen) and cleaned using a RNeasy® Mini Kit (Qiagen). One μg of the RNA was used to generate cDNA using an iScript cDNA synthesis kit (Bio-Rad). qRT-PCR analyses were carried out with CFX96 Real-Time System (Bio-rad), using Maxima SYBR Green qPCR Master Mix (Thermo Fisher Scientific Co.), and relative expression values were calculated using the Δ-Ct-method. A complete list of the primers used for RT-PCR is presented in Supplementary Table S1.

### Vacuum-infiltration of germinated seeds

To perform vacuum-infiltration of desired constructs, around 30-50 seeds of *P. fendleri* were surface sterilized with 95% ethanol for 1 min and washed with sterilized water three times. To promote uniform seed germination provided 3 mM GA3 treatment for 2 hours and washed the seeds with sterilized water three times and transferred them to wet blotting papers and kept them in dark for 2 days to germinate. Next day, started preparation for the agrobacterium culture containing pTRV1 and pTRV2-target plasmids. The primary 5-mL agrobacterium culture containing pTRV1 and pTRV2-target plasmids was grown overnight at 28°C with appropriate antibiotics. The culture was inoculated into 10 mL of Luria-Bertani medium containing antibiotics, the culture was grown 4-5 hours at 28°C shaker, to around OD_600_ of 0.3-0.5. 10 mL culture was made in 50 mL falcon tubes by mixing 5 mL pTRV1 and 5 mL pTRV2-target with supplements (Zhang et al., 2017) (100mM Cysteine, 50 μL Tween-20, and freshly prepared 100mM Acetosyringone). Around 50-60 germinated seeds were added to the 10 mL culture, make sure that all the seeds dipped in culture and vacuum infiltrate for 5 min at 15 psi. The seeds were co-cultivated with in agro in dark on MS plate or wet filter paper (1/2 MS media) for one day. The co-cultivated seeds were washed with sterilized water and grown in pots in a growth room of 16/8 hrs light/dark cycle, light of 240-250 μmol/m^2^/sec at 23°C.

### Lipid quantification by GC-FID

Total lipids was extracted from mature seeds using hexane-isopropanol method (Hara and Radin, 1978) with some modification as described in. An aliquot of total lipid was converted to fatty acid methyl esters (FAME) for fatty acid composition and quantification by gas chromatography with flame ionization detection (GC-FID) (Karki and Bates, 2018).

## Acknowledgements

This work supported by the United States Department of Agriculture National Institute of Food and Agriculture grants nos. 2020-67013-30899 and 2023-67013-39022.

## Author contributions

P.D.B. and A.A. conceived the research. A.A. designed and executed the experiments. Both authors wrote the article.

