## Supplemental for "Self-incompatibility based functional genomics for rapid phenotypic characterization of seed metabolism genes"

**Table S1. List of primers used in this study.**

| Primer name | Sequence (5'-3) | Purpose |
|---|---|---|
| pTRV2_F | GTTACTCAAGGAAGCACGATGAGC | Colony PCR |
| pTRV2_R | CGTAGTTTAATGTCTTCGGGACATGC | Colony PCR |
| PfePDS_F1 | CGCGAATTCGAGCGCGTGACAGACGA | Cloning |
| PfePDS_R1 | CGCTCTAGATGCACAATAGACTGTGAGCAGA | Cloning |
| PfeSRK_F1 | CGCGAATTCGGTGAGGTCTTCGAGCTTGG | Cloning |
| PfeSRK_R1 | CGCGGATCCTTCATCTGTGGAAGTAAAGTATCA | Cloning |
| PfeSCR_F | CGGATCCATGAAAATTGGTACTTTCTTCATAGTTTC | Cloning |
| PfeSCR_R | CGCGGTACCGGTCTCGATTTAGTAGCAGCAAAC | Cloning |
| PfeFAE1-F | CGCGGTACCCATAAAGCTCCTTTACCATTACATCC | Cloning |
| PfeFAE1-R | CGCGAGCTCATAACCATGGCTGATAGAGAAGG | Cloning |
| PfeFAH12_F | CGCGAATTCGTTACCCCCTCTTCCAAGAAA | Cloning |
| PfeFAH12_R | CGCGGATCCTGTATATCTGGAGGCGTTCTCG | Cloning |
| Pfe18S_F | GAGAAACGGCTACCACATCCA | qPCR |
| Pfe18S_R | CCGTGTCAGGATTGGGTAATTT | qPCR |
| PfeSRK_qF | GGTGAGGTCTTCGAGCTTGG | qPCR/Semi-qPCR |
| PfeSRK_qR | TTCATCTGTGGAAGTAAAGTATCA | qPCR/Semi-qPCR |
| PfeSCR_qF | ATGAAAATTGGTACTTTCTTCATAGTTTC | Semi-qPCR |
| PfeSCR_qR | GGTCTCGATTTAGTAGCAGCAAAC | Semi-qPCR |

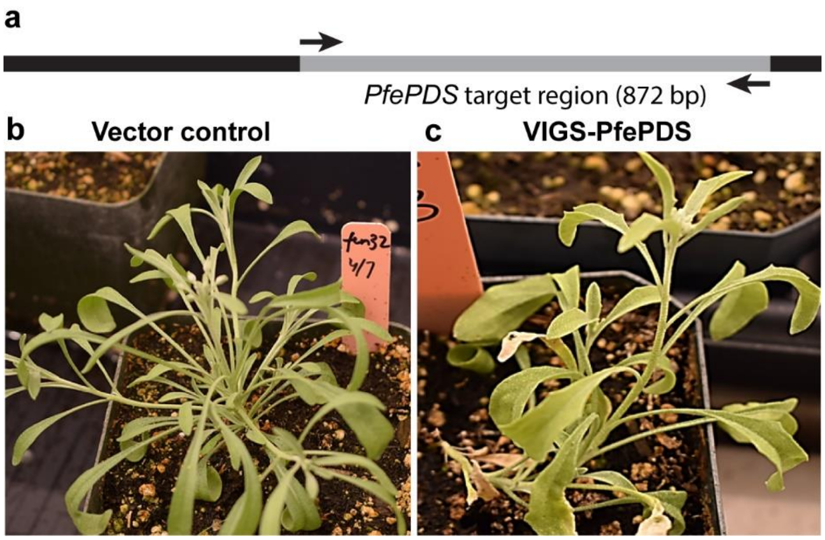


**Figure S1. Agrobacterium leaf-infiltration of *pTRV2-PfePDS construct.*** (a) Schematic representation of the *PfePDS* target region (872 bp). (b) Leaf-infiltrated vector control (pTRV2) plants show no photobleaching phenotype. (c) Leaf-infiltrated pTRV2-PfePDS plants show partial photobleaching phenotype (light green leaves). Leaf-infiltration was performed at the four leaf stage following.

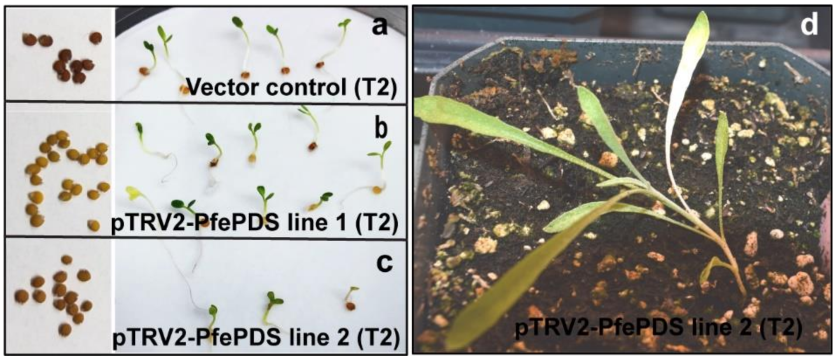


**Figure S2.** ***pTRV2-PfePDS VIGS plants shows heritable photobleaching phenotype***. (a) Vector control T2 seeds, dark brown in color and show no photobleaching phenotype after germination, (b), (c). pTRV2-PfePDS VIGS T2 seeds are light brown in color and some seeds after germination show heritable photobleaching phenotype, (d). Partial photobleaching phenotype in pTRV2-PfePDS T2 plants.

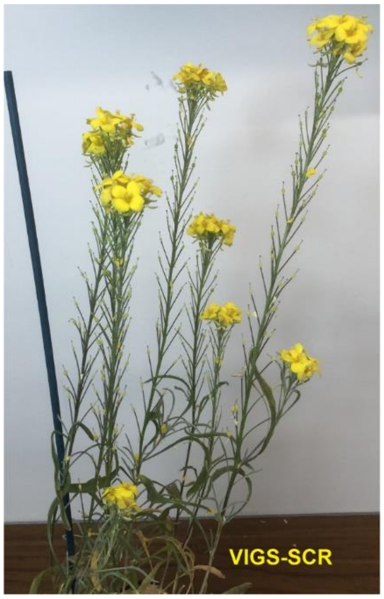


**Figure S3. Virus induced gene silencing of *P. fendleri* SCR produced no seed without out-cross.**
